# Genomic imprinting mediates dosage compensation in a young plant XY system

**DOI:** 10.1101/179044

**Authors:** Aline Muyle, Niklaus Zemp, Cécile Fruchard, Radim Cegan, Jan Vrana, Clothilde Deschamps, Raquel Tavares, Roman Hobza, Franck Picard, Alex Widmer, Gabriel AB Marais

## Abstract

This preprint has been reviewed and recommended by Peer Community In Evolutionary Biology (http://dx.doi.org/10.24072/pci.evolbiol.100044).

Sex chromosomes have repeatedly evolved from a pair of autosomes^1^. Consequently, X and Y chromosomes initially have similar gene content, but ongoing Y degeneration leads to reduced Y gene expression and eventual Y gene loss. The resulting imbalance in gene expression between Y genes and the rest of the genome is expected to reduce male fitness, especially when protein networks have components from both autosomes and sex chromosomes. A diverse set of dosage compensating mechanisms that alleviates these negative effects has been described in animals^2–4^. However, the early steps in the evolution of dosage compensation remain unknown and dosage compensation is poorly understood in plants^5^. Here we show a novel dosage compensation mechanism in the evolutionarily young XY sex determination system of the plant *Silene latifolia*. Genomic imprinting results in higher expression from the maternal X chromosome in both males and females. This compensates for reduced Y expression in males but results in X overexpression in females and may be detrimental. It could represent a transient early stage in the evolution of dosage compensation. Our finding has striking resemblance to the first stage proposed by Ohno for the evolution of X inactivation in mammals.

---

In *Drosophila*, the X chromosome is upregulated specifically in males, resulting in complete dosage compensation through both ancestral expression recovery in males and equal expression between the sexes (hereafter sex equality)^6^. In *Caenorhabditis elegans*, both X chromosomes are downregulated in XX hermaphrodites resulting in sex equality, but only a few genes have their X expression doubled for ancestral expression recovery^7^. In placental mammals, including humans, one X chromosome is randomly inactivated in XX females, resulting in sex equality but without recovering the ancestral expression of sex chromosomes, except for a few dosage-sensitive genes whose X expression was doubled in both sexes^8–12^. In the marsupials, the paternal X chromosome is consistently inactivated in XX females^13^. Differential expression that depends on the parent of origin is known as genomic imprinting^14^, and this mechanism also operates in the mouse placenta^15^.

Despite the plethora of studies on gene expression on sex chromosomes, it is not yet clear if genomic imprinting is commonly involved in the early steps of dosage compensation evolution. In a seminal work, Ohno hypothesized a two-step process for the evolution of dosage compensation^16^. In the first step, expression from the X is doubled, thereby mediating the recovery of ancestral expression in XY males. Second, the resulting overexpression in XX females selects for X inactivation. This scenario is consistent with the fact that sexual selection is often stronger on males than on females. Under this scenario, selection on XY males to upregulate their single X chromosome should be stronger than selection on females, leading to overexpression in females until a second correcting mechanism evolves^3^. However, in order to understand these early steps of dosage compensation evolution, species with young sex chromosomes must be studied.

The plant *Silene latifolia* is an ideal model to study early steps of sex chromosome evolution thanks to its pair of X/Y chromosomes that evolved ∼4 Mya^17^. Dosage compensation is poorly understood in plants^5^. Thus far only sex equality has been studied. Equal expression levels were observed for males and females for some genes despite Y expression degeneration^18–23^. However, the mechanisms through which sex equality is achieved – and whether ancestral expression is recovered in *S. latifolia* males – remain unknown. To address these questions, we have developed an approach relying on (i) the use of an outgroup without sex chromosomes as an ancestral autosomal reference^5^ in order to determine whether X chromosome expression increased or decreased in *S. latifolia*, (ii) the application of methods to study allele-specific expression while correcting for reference mapping bias^5^, and (iii) a statistical framework to quantify dosage compensation^5^.

Because only ∼25% of the large and highly repetitive *S. latifolia* genome has been assembled so far^23^, we used an RNA-seq approach based on the sequencing of a cross (parents and a few offspring of each sex), to infer sex-linked contigs (i.e. contigs located on the non-recombining region of the sex chromosome pair)^24^. X/Y contigs show both X and Y expression, while X-hemizygous contigs are X-linked contigs without Y allele expression. We made inferences separately for three tissues: flower buds, seedlings and leaves (Supplementary Table S2). Results are consistent across tissues and flower buds and leaves are shown in Supplementary Materials. In seedlings, ∼1100 sex-linked contigs were inferred. Among these, 15% of contigs with significant expression differences between males and females were removed for further analyses (Supplementary Table S2 and Materials and Methods). These are likely involved in sex-specific functions and are not expected to be dosage compensated^25^. This was done as a usual procedure for studying dosage compensation, however the resulting trends and significance levels are not affected. About half of the non sex-biased sex-linked contigs could be validated by independent data using three sources: literature, a genetic map and sequence data from Y flow-sorted chromosomes (see Supplementary Table S2 and Materials and Methods). X-hemizygous contigs are more difficult to identify than X/Y contigs using an RNA-seq approach (see Supplementary Text S1). This explains conflicting earlier results on dosage compensation in *S. latifolia^5^.* A study using genomic data (i.e. not affected by the aforementioned ascertainment bias) found sex-equality in approximately half of the studied X-hemizygous genes^23^. In our set of X-hemizygous contigs, no evidence for dosage compensation was found (Supplementary Text S1), in agreement with previous work relying on an RNA-seq approach^18,22^. This could be due to an over-representation of dosage insensitive genes in our set of X-hemizygous contigs (Supplementary Text S1).

We estimated paternal and maternal allele expression levels in males and females for sex-linked and autosomal contigs in *S. latifolia* after correcting for reference mapping bias (Materials and Methods). We then compared these allelic expression levels to one or two closely related outgroups without sex chromosomes in order to polarise expression changes in *S. latifolia.* For autosomal contigs, expression levels did not differ between *S. latifolia* and the outgroups (Figure 1). This is due to the close relatedness of the outgroups (∼5My, Supplementary Figure S1), and validates their use as a reference for ancestral expression levels. We used the ratio of Y over X expression levels in *S. latifolia* males as a proxy for Y degeneration and then grouped contigs on this basis. As expression of the Y allele decreased (paternal allele in blue in Figure 1), expression of the corresponding X allele in males increased (maternal allele in red in Figure 1). This is the first direct evidence for ancestral expression recovery in *S. latifolia*, i.e. ancestral expression levels are reestablished in males despite Y expression degeneration. In females, expression of the maternal X allele also increased with Y degeneration (gray bars in Figure 1), similarly to the maternal X allele in males. The paternal X alleles in females, however, maintained ancestral expression levels, regardless of Y degeneration (black bars in Figure 1). Consequently, sex equality is not achieved in *S. latifolia* due to upregulation of sex-linked genes in females compared to ancestral expression levels. These results were confirmed in two other tissues and when analysing independently validated contigs only (although statistical power is sometimes lacking due to the limited number of validated contigs, Supplementary Figures S2-S7).

**Figure 1:**
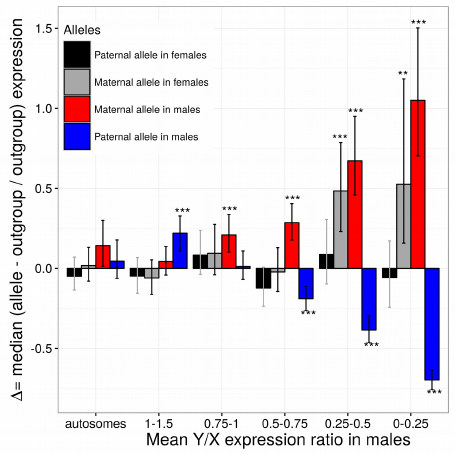
Normalised difference (hereafter Δ) in allelic expression levels between *S. latifolia* and the outgroup without sex chromosomes *S. vulgaris*, in autosomal and sex-linked contigs for the seedling tissue. If Δ is lower, higher or equal to zero, then expression in *S. latifolia* is respectively lower, higher or equal to the outgroup (See Materials and Methods for details). For all contig categories, A was compared to zero using a Wilcoxon test. The median Δ, confidence intervals and p-values adjusted for multiple testing using a Benjamini and Hochberg correction are shown (***: p-value < 0.001; **: p-value < 0.01, *: p-value < 0.05). Allelic expression at SNP positions was averaged for each contig separately and the Y/X ratio was used as a proxy for Y degeneration to group contigs. Contigs with sex-biased expression were removed, as well as contigs with Y/X expression ratios above 1.5. Sample sizes for the different contig categories are: autosomal: 200; 1-1.5:148; 0.75-1:139; 0.5-0.75:160; 0.25-0.5:114; 0-0.25:79 (we randomly selected 200 autosomal contigs to ensure similar statistical power among gene categories).

Upregulation of the maternal X allele both in males and females of *S. latifolia* (Figure 1 and Supplementary Figures S2-S7) establishes a role for genomic imprinting in dosage compensation. In order to statistically test this inference at the SNP level, we used a linear regression model with mixed effects (Materials and Methods). Outgroup species were used as a reference and expression levels in *S. latifolia* were then analyzed while accounting for the variability due to contigs and individuals. The joint effect of the parental origin and the degeneration level was estimated, which allowed computing expression differences between maternal and paternal alleles in females for different Y/X degeneration categories (Figure 2). Maternal and paternal alleles of autosomal SNPs were similarly expressed in females, indicating a global absence of genomic imprinting for these SNPs. However, for X/Y SNPs, the difference between the maternal and paternal X in females increased with Y degeneration. These results were confirmed in two other tissues and when analysing independently validated contigs only (although statistical power is sometimes lacking due to the limited number of validated contigs, Supplementary Figures S8-S13).

**Figure 2:**
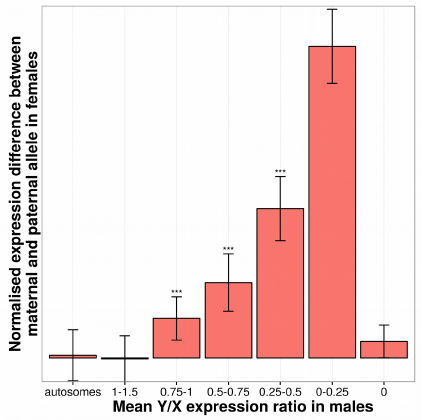
Normalised expression difference between maternal and paternal alleles in *S. latifolia* females in autosomal and sex-linked SNPs in the seedling tissue. The Y axis unit is the normalised allelic read count difference in log scale. A linear regression model with mixed effects was used to estimate the normalised difference between the effect of paternal and maternal origin of alleles in interaction with the contig status (autosomal or sex-linked with various levels of Y degeneration), while accounting for inter-contig and inter-individual variability (see Materials and Methods for details). The analysis is SNP-wise and reveals consistent patterns across SNPs. See Fig. 1 legend for sample sizes for the different contig categories and statistical significance symbols.

Previous studies that showed sex equality in *S. latifolia* could have been explained by simple buffering mechanisms, where one copy of a gene is expressed at a higher level when haploid than when diploid, due to higher availability of the cell machinery or adjustments in gene expression networks^23,26,27^. However, the upregulation of the X chromosome we reveal here in *S. latifolia* males cannot be explained by buffering mechanisms alone, as the maternal X in females would otherwise not be upregulated. Instead, our findings indicate that a specific dosage compensation mechanism relying on genomic imprinting has evolved in *S. latifolia.* This apparent convergent evolution with marsupials is mediated by different mechanisms (in marsupials the paternal X is inactivated^13^, while in *S. latifolia* the maternal X is upregulated).

An exciting challenge ahead will be to understand how upregulation of the maternal X is achieved in *S. latifolia* males and females at the molecular level. Chromosome staining suggests that DNA methylation is involved. Indeed, one arm of one of the two X chromosomes in females was shypomethylated, as well as the same arm of the single X in males^28^ (Figure 3 and Supplementary Figure S14). Based on our results, we hypothesize that the hypomethylated X chromosome corresponds to the maternal, upregulated X. Unfortunately, parental origin of the X chromosomes was not established in this study^28^. It would be of interest in the future to study DNA methylation patterns in *S. latifolia* paternal and maternal X chromosomes, along with the homologous pair of autosomes in a closely related species without sex chromosomes. The methylation pattern observed by chromosome staining suggests that dosage compensation in *S. latifolia* could be a chromosome arm-wide phenomenon. To test this hypothesis with expression data, positions of genes along the X chromosome remain to be elucidated.

**Figure 3:**
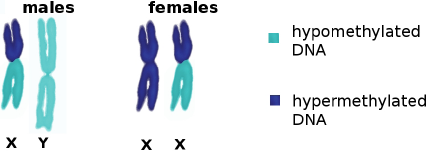
Illustration of DNA methylation staining results in *S. latifolia* from Siroky et al. ^28^. See Supplementary Figure S14 for the original Figure. One arm of one of the two X chromosomes in females was hypomethylated, as well as the same arm of the single X in males.

Our study is the first to establish female upregulation of the X chromosome compared to autosomes, as predicted by Ohno. An earlier report in *Tribolium castaneum was* later shown to be due to biases from inclusion of gonads in whole body extracts^4^. X overexpression in females may be deleterious. Its occurence suggests that reduced expression of sex-linked genes in males is more deleterious than overexpression in females. This potentially suboptimal situation may be transitory and a consequence of the young age of *S. latifolia* sex chromosomes. Sex equality may evolve at a later stage, following the evolutionary path trajectory originally proposed by Ohno for placental mammals^16^.

## Methods

### Sequence data and inference of sex-linkage

RNA-seq data was generated in *S. latifolia* for a cross (parents and progeny) for three tissues (seedlings, leaves and flower buds) and analysed using the SEX-DETector pipeline^24^. RNA-seq data was also generated for two outgroup species (*S. viscosa* and *S. vulgaris).* Reference mapping bias was corrected using the program GSNAP^29^. Inferences of sex-linked contigs were validated using three sources of information (literature, a genetic map and flow-sorted Y chromosome sequences). See Supplementary Text S2 for details.

### Allelic expression levels

Contigwise autosomal, X, Y, X+X and X+Y normalised allelic expression levels were computed by summing read numbers for each X-linked or Y-linked alleles for filtered SNPs of the contigs (Supplementary Text S2) for each individual separately and then normalised using the library size and the number of studied sex-linked SNPs in the contig:

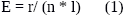

With E = normalised expression level for a given individual, r = sum of total read counts, n = number of studied SNPs, l = library size of the individual (number of mapped reads). Allelic expression levels were then averaged among individuals for each contig. In order to make *S. latifolia* expression levels comparable to *S. viscosa* and *S. vulgaris*, *S. vicosa* and *S. vulgaris* expression levels were estimated using only the filtered SNP positions used in *S. latifolia*. Normalised expression levels computed as explained in equation (1) in the two outgroups were then averaged together for leaves and flower buds as expression levels are highly correlated (R^2^ 0.7 and 0.5 for flower buds and leaves respectively and p-value < 2.10^-6^ in both cases). Averaging expression levels between the two outgroups allows to get closer to the ancestral autosomal expression level.

### Sex-biased expression

Sex-biased contigs were inferred as in Zemp et al^30^. See Supplementary Text S2 for more detail.

### Expression divergence between *S. latifolia* and the two outgroups at the contig level

The normalised difference in allelic expression between *S. latifolia* and the two outgroups (hereafter Δ) was computed in order to study how sex chromosome expression levels evolved in *S. latifolia* compared to autosomal expression levels in the two outroups: Δ is equal to zero if *S. latifolia* and the outgroups have equal expression levels, Δ is positive if *S. latifolia* has higher expression levels compared to the outgroups and Δ is negative otherwise:

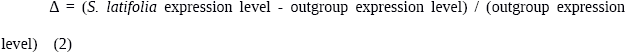

Sex-linked contigs were grouped by categories of degeneration level using the average Y over X expression ratio in males. 200 autosomal contigs were randomly selected in order to have similar statistical power among gene categories. Δ values for each allele (maternal and paternal in males and females) and each gene category were compared to zero using a Wilcoxon test. P-values were corrected for multiple testing using a Benjamini and Hochberg correction. The estimated median Δ, confidence intervals and adjusted p-values were then used to plot Figure 1 and Supplementary Figures S2 to S7.

### Expression differences between maternal and paternal alleles at the SNP level

Maternal and paternal alleles expression were compared in *S. latifolia* for autosomal and sex-linked SNPs. In order to deal with the difference in numbers of autosomal versus sex-linked contigs (Supplementary Table S2), 200 autosomal contigs were randomly selected in order to keep comparable powers of detection. Allelic expression levels in *S. latifolia* for each individual at every SNP position were analysed using a linear regression model with mixed effects with the R package lme4. We assumed a normal distribution of the read count data after log transformation. In order to account for inter-individual and inter-contig variability, a random “individual” and a random “contig” effect were included in the model. The aim of this modeling framework was to estimate the joint effect of the chromosomal origin of alleles (paternal or maternal in males or females) and the status of the gene (autosomal or sex-linked with various levels of Y degeneration defined by the average Y over X expression ratio in males). Two fixed effects with interaction were therefore considered in the model, see equation (3). In order to estimate the changes in sex-linked gene expression levels since the evolution of sex chromosomes, we used the average of the two outgroup expression levels as a reference (offset) for every SNP position, divided by two in order to be comparable to *S. latifolia* allelic expression levels.

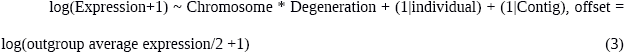

All effects of the model (fixed or random) were proved highly significant (p-values < 2.2.10^-16^) using comparison of the fit of model (3) to simpler nested models (removing one effect at a time in model (3)). In order to statistically test whether there was a difference between the effects of paternal and maternal alleles in females in different degeneration categories we used the contrasts provided by the lmerTest package in R. This strategy provided estimates, confidence intervals and p-values of the difference between the two effects of paternal and maternal origin in females in interaction with degeneration levels, while normalising by the expression of the two outgroups. Moreover, the presence of random effects allows to account for inter-individual and inter-contig variability. Finally, p-values were corrected for multiple testing using a Benjamini and Hochberg correction. These values were used to plot Figure 2 and Supplementary Figures S8 to S13.

## Data Availability

The new sequence data presented here can be downloaded from the European Nucleotide Archive (ENA) under accession number PRJEB24933.

## Supplementary Materials

Supplementary Information includes Supplementary Texts S1-S2, Supplementary Figures S1-S14 and Supplementary Tables S1-S3.

## Acknowledgments

This project was supported through an ANR grant ANR-14-CE19-0021-01 to G.A.B.M, SNSF grant 160123 to A.W and Czech Science Agency grant number 16-08698S to R.H. We thank Brandon Gaut, Tatiana Giraud and Judith Mank for comments on the manuscript.

## Author contributions

Aline Muyle, Niklaus Zemp, Alex Widmer and Gabriel Marais conceived the study and experimental design. Niklaus Zemp and Alex Widmer prepared and sequenced the plant material. Aline Muyle ran SEX-DETector on the RNA-seq datasets for the three tissues, analysed the data, prepared Tables and Figures and wrote the Supplementary Material with inputs from other authors. Niklaus Zemp generated the X chromosome genetic map (with help from Aline Muyle for the mapping and genotyping part). Radim Cegan, Jan Vrana and Roman Hobza did the Y chromosome flow cytometry sorting and sequencing. Clothilde Deschamps did the first assembly of the sorted Y chromosome and improved it with RNA-seq data with the help of Cecile Fruchard. Aline Muyle did the blasts to validate the inferences of SEX-DETector. Raquel Tavares did the GO term analysis. Aline Muyle and Frank Picard did the statistical analyses of the data. Gabriel Marais and Aline Muyle wrote the main text of the manuscript with inputs from other authors.

## Author information

The authors declare no competing interests.

Correspondence and requests for materials should be addressed to Aline Muyle.

## Supplementary Materials

### Supplementary Text S1: Dosage compensation in X-hemizygous genes

The first papers on dosage compensation in *S. latifolia* were contradictory because they focused on different gene sets. Muyle et al.^1^ focused on X/Y gene pairs while other papers focused on X-hemizygous genes^2,3^. However, the X-hemizygous gene sets returned by the RNA-seq approach used in those papers is less reliable than the X/Y gene sets^4^. A gene might be inferred as X-hemizygous simply because the – still functional – Y copy is not expressed in the tissue sampled for RNA-seq. In *S. latifolia*, X-hemizygous genes tend to be less expressed than X/Y genes and are less likely to be detected by segregation analysis as efficient SNP calling requires a certain read depth, see^4^. Moreover, X-hemizygous genes are inferred from X polymorphisms while X/Y genes can be detected both with X and X/Y polymorphisms, which are more numerous. Another inherent bias to X-hemizygous contig inference comes from the assembly step. If the X and the Y copy are too divergent to be assembled together, the X contig will be wrongly inferred as X-hemizygous because Y alleles will be absent from the contig (this bias was at least partly corrected in the analyses presented here, see Material and Method section 5.1). The inferences of X-hemizygous genes using the RNA-seq approach (including SEX-DETector) imply a higher rate of both false positives and false negatives than those for X/Y gene pairs. In Papadopoulos et al.^5^, 25% of the X/Y chromosomes were sequenced using a genomic approach. A much higher fraction of X-hemizygous genes was found than in previous RNA-seq papers^2,3^. Papadopoulos et al.^5^ did find evidence for dosage compensation in approximately half of X-hemizygous genes (see their figure 3D). Due to limitations of the RNA-seq approach in inferring X-hemizygous genes, results on X-hemizygous contigs are analysed separately here.

Poor dosage compensation of X-hemizygous contigs compared to X/Y contigs with high Y degeneration was observed across all tissues (Supplementary Figures 2 to 7). Also, the parental origin of the X chromosome has limited to no effect on female X expression levels for X-hemizygous contigs, unlike X/Y contigs (Supplementary Figures 8 to 13). A reason that could explain such a different pattern for X-hemizygous genes compared to X/Y genes is the possible dosage insensitivity of X-hemizygous genes. X-hemizygous genes could have lost their Y copy because dosage was not important for them and selection neither slowed down the loss of the Y copy nor selected for dosage compensation when degeneration inevitably occurred^6^. A well described characteristic of dosage sensitive genes is that they tend to code proteins involved in large complexes^7^. Gene Ontology was studied using the Blast2GO PRO version 2.7.2^30^ as in^8^. Using the GO-term analysis, our set of X-hemizygous contigs were found to be significantly depleted in ribosomal protein coding genes compared to autosomal genes (p-value 1.3.10^-4^), which is consistent with the global dosage insensitivity of X-hemizygous genes in *S. latifolia*. This depletion in large protein complexes was not found when comparing X/Y genes to autosomal genes.

### 1) Plant material and sequencing

#### 1.1) RNA-seq Illumina data

RNA-seq data from previous studies were used (the GEO database GEO Series GSE35563, European Nucleotide Archive PRJEB14171), it included flower buds and leaf tissues from individuals of a cross in *S. latifolia* as well as individuals in *S. vulgaris*. In addition to these preexisting data, RNA-seq reads were generated in a comparable way for seedlings of a controlled cross using the same parents in *S. latifolia*, four males and four females were sampled (Seed_lati_female_1, Seed_lati_female_2, Seed_lati_female_3, Seed_lati_female_4, Seed_lati_male_1, Seed_lati_male_2, Seed_lati_male_3 and Seed_lati_male_4). Seedlings were also sequenced for *S. vulgaris* (Seed_vulg_herm_1, Seed_vulg_herm_2, Seed_vulg_herm_3 and Seed_vulg_herm_4). Seedlings were grown in a temperature controlled climate chamber in Eschikon (Switzerland) using the same conditions as in^8^. The *S. latifolia* and *S. vulgaris* seedlings were collected without roots at the four-leaf stage. The sexing of the *S. latifolia* seedlings was done using Y specific markers^9^ that were amplified with the direct PCAR KAPA3G Plant PCR Kit (however male number 3 was later shown to be a female). High quality RNA (RIN > 8.5) was extracted using the total RNA mini kit from Geneaid. Twelve RNA-seq libraries were produced using the Truseq kit v2 from Illimina. Libraries were tagged individually and sequenced in two Illumina Hiseq 2000 channels at the D-BSSE (ETH Zürich, Switzerland) using 100 bp paired-end read protocol.

*S.viscosa* seeds we received from botanical gardens or collected in the wild by Bohuslav Janousek and grown under controlled conditions in a greenhouse in Eschikon (Switzerland) and Lyon (France). Similarly to^8^, flower buds after removing the calyx and leaves were collected. Total RNA were extracted through the Spectrum Plant Total RNA kit (Sigma, Inc., USA) following the manufacturer’s protocol and treated with a DNAse. Libraries were prepared with the TruSeq RNA sample Preparation v2 kit (Illumina Inc., USA). Each 2 nM cDNA library was sequenced using a paired-end protocol on a HiSeq2000 sequencer. Demultiplexing was performed using CASAVA 1.8.1 (Illumina) to produce paired sequence files containing reads for each sample in Illumina FASTQ format. RNA extraction and sequencing were done by the sequencing platform in the AGAP laboratory, Montpellier, France (http://umr-agap.cirad.fr/).

A female individual from an interspecific *S. latifolia* cross (C1_37) was back crossed with a male from an 11 generation inbred line (U10_49). The offspring (hereafter called BC1 individuals) were grown under controlled conditions in a greenhouse in Eschikon (Switzerland). High quality RNA from flower buds as described in^10^ was extracted from 48 BC1 individuals (35 females and 13 males). 48 RNA-seq libraries were produced using the Truseq kit v2 from Illimina with a median insert size of about 200 bp. Individuals were tagged separately and sequenced in four Illumna Hiseq 2000 channels at the D-BSSE (ETH Zürich, Switzerland) using 100bp paired-end read protocol. The parents used for this back cross had previously been sequenced in a similar way^1,8^.

#### 1.2) DNA-seq data from filtered Y chromosome

Y chromosome DNA was isolated using flow cytometry. The samples for flow cytometric experiments were prepared from root tips according to^11^ with modifications. Seeds of *S. latifolia* were germinated in a petri dish immersed in water at 25°C for 2 days until optimal length of roots was achieved (1 cm). The root cells were synchronized by treatment with 2mM hydroxyurea at 25°C for 18h. Accumulation of metaphases was achieved using 2.5μM oryzalin. Approximately 200 root tips were necessary to prepare 1ml of sample. The chromosomes were released from the root tips by mechanical homogenization using a Polytron PT1200 homogenizer (Kinematica AG, Littau, Switzerland) at 18,000rpm for 13 s. The crude suspension was filtered and stained with DAPI (2μg/ml). All flow cytometric experiments were performed on FACSAria II SORP flow cytometer (BD Biosciences, San José, Calif., USA). Isolated Y chromosomes were sequenced with 2×100bp PE Illumina HiSeq.

#### 1.3) RNA-seq PacBio data

Plants from an 11 generation inbred line were grown under controlled conditions in a greenhouse in Eschikon (Switzerland). One male (U11_02) was randomly selected. High quality RNA (RIN > 7.5) were extracted using the total RNA mini kit of Geneaid from very small flower buds, small and large flower buds, flowers before anthesis without calyces, rosette leaves, seedlings (4 leaves stage) and pollen. RNA of the different tissues was equally pooled and cDNA was produced using the Clontech SMARTer Kit. The cDNA pool was then normalized using a duplex specific endonuclease of the Evrogen TRIMMER kit. Two ranges were selected (1- 1.3 kb and 1.2 -2 kb) using the Pippin Prep (Sage Science). Two SMRTbell libraries were prepared using the C2 Pacific Biosciences (PacBio) chemistry and sequenced with two SMRT Cells runs on a PacBio RS II at the Functional Genomic Center Zurich (FGCZ).

#### 1.4) RNA-seq 454 data

Previously generated 454 data was used^8,12^.

### 2) Reference trancriptome assembly

The same reference transcriptome as in Muyle et al.^12^ and Zemp et al.^8^ was used.

### 3) Inference of sex-linked contigs

Autosomal and sex-linked contigs were inferred as in Muyle et al.^12^ and Zemp et al.^8^. Illumina reads from the individuals of the cross were mapped onto the assembly using BWA^13^ version 0.6.2 with the following parameters: bwa aln -n 5 and bwa sampe. The libraries were then merged using SAMTOOLS version 0.1.18^14^. The obtained alignments were locally realigned using GATK IndelRealigner^15^ and were analysed using reads2snps^16^ version 3.0 with the following parameters: -fis 0 -model M2 -output_genotype best -multi_alleles acc -min_coverage 3 -par false. This allowed to genotype individuals at each loci while allowing for biases in allele expression, and without cleaning for paralogous SNPs. Indeed, X/Y SNPs tend to be filtered out by paraclean, a program which removes paralogous positions^17^. A second run of genotyping was done with paraclean in order to later remove paralogous SNPs from autosomal contigs only. SEX-DETector^12^ was then used to infer contig segregation types after estimation of parameters using an SEM algorithm. Contig posterior segregation type probabilities were filtered to be higher than 0.8. Because the parents were not sequenced for the leaf and seedling datasets, SEX-DETector was run using the flower bud data for the parents.

### 4) Reference mapping bias correction

In order to avoid biases towards the reference allele in expression level estimates, a second mapping was done using the program GSNAP^18^ with SNP tolerant mapping option. A GSNAP SNP file was generated by home-made perl scripts using the SEX-DETector SNP detail output file. Shortly, for each polymorphic position of all contigs, the most probable posterior SNP type was used to extract the possible alleles and write them to the GSNAP SNP file. This way, reference mapping bias was corrected for both sex-linked and autosomal contigs. Only uniquely mapped and concordant paired reads were kept after this. See Supplementary Table S1 for percentage of mapped reads. SEX-DETector was run a second time on this new mapping and the new inferences were used afterwards for all analyses (see Supplementary Table S2 for inference results).

### 5) Validation of sex-linked contigs

#### 5.1) Detection of false X-hemizygous contigs

Erroneous inference of X-hemizygous contig can be due to a true X/Y gene which X and Y copies were assembled into different contigs. In order to detect such cases, X-hemizygous contigs were blasted^19^ with parameter -e 1E-5 against RNA-seq contigs that have male-limited expression (see section 7 below for how male-limited contigs were inferred). These cases were removed from the analyses presented here.

#### 5.2) Validation using data from literature

A few sex-linked and autosomal genes in *S. latifolia* have already been described in the literature (see Supplementary Table S3).

#### 5.3) Validation using a genetic map

A genetic map was built and contigs from the X linkage group were used to validate SEX-DETector inferences. RNA-seq reads from the flower bud *S. latifolia* full-sib cross (hereafter CP) and backcross (hereafter BC1) were mapped against the reference transcriptome using BWA^13^ with a maximum number of mismatch equal to 5. Libraries were merged and realigned using GATK^15^ and SNPs were analysed using reads2snps^16^. Using a customized perl script, SNP genotypes from the parents and the offspring as well as the associated posterior probabilities were extracted from the reads2snps output file. Only SNPs with a reads2snps posterior genotyping probability higher than 0.8 were kept for further analyses. Then, only informative SNPs were kept: both parents had to be homozygous and different between father LEUK144-3 and mother U10_37 in a first generation backcross population design (BC1) and at least one allele had to be different between mother C1_37 and father U10_49 in the cross-pollinator (CP). Filtered SNPs were then converted into a JoinMap format using a customized R script. If more than one informative SNP per contig was present, the SNP was used with less segregation distortion and less missing values. This led to 8,023 BC1 and 16,243 CP markers.

Loci with more than 10 *%* missing values were excluded, resulting in 7,951 BC1 and 15,118 CP markers. Linkage groups were identified using the default setting of JoinMap 4.1^20^. Robustness of the assignment of the linkage groups was tested using LepMap^21^. Blasting the contigs against known sex-linked genes allowed the identification of the X chromosome linkage group. Contigs could not be ordered along the linkage groups due to the too limited number of individuals that prevented the convergence of contig order. However, contigs were reliably attributed to linkage groups.

#### 5.4) Validation using isolated Y chromosome DNA-seq data

Filtered Y chromosome DNA-seq reads were filtered for quality and Illumina adapters were removed using the ea-utils FASTQ processing utilities^22^. The optimal kmer value for assembly was searched using KmerGenie^23^. Filtered reads were assembled using soapdenovo2^24^ with kmer=49, as suggested by KmerGenie. The obtained assembly was highly fragmented, therefore RNA-seq data was used to join, order and orient the genomic fragments with L_RNA_scaffolder^25^. The following RNA-seq reads were used (see section 1): one sample of male flower buds sequenced by 454, 6 samples of male flower buds sequenced by Illumina paired-end, 4 samples of male leaves sequenced by Illumina paired-end and one sample of male pooled tissues sequenced by PacBio. The genomic assembly was successively scaffolded with L_RNA_scaffolder using RNA-seq samples one after the other, first 454 samples then Illumina and finally PacBio. The obtained contigs were filtered to be longer than 200pb.

#### 5.5) Set of validated sex-linked and autosomal contigs

The three sources of data (litterature, genetic map and filtered Y sequence data) were compared to SEX-DETector inferred sex-linked RNA-seq contigs using BLAST^19^ with parameter -e 1E-5. Blasts were filtered for having a percentage of identity over 90%, an alignment length over 100bp and were manually checked. If a sex-linked RNA-seq contig blasted against a sequence from one of the three data sources (literature, X genetic map or filtered Y DNA-seq) it was then considered as validated. See Supplementary Table S2 for numbers of validated sex-linked contigs.

### 6) Expression level estimates

#### 6.1) whole contig expression levels

Whole contig mean expression levels were obtained for each individual using GATK DepthOfCoverage^15^ as the sum of every position coverage, divided by the length of the contig. Normalised expression levels, in RPKM^26^, were then computed for each individual by dividing by the value by the library size of the individual (total number of mapped reads), accounting for different depths of coverage among individuals. Whole contig mean male and female expression levels were then computed by averaging male and female individuals for each contig.

#### 6.2) Allelic expression levels filtering

In order to study separately X and Y allele expression levels in males and females, expression levels were studied at the SNP level. In *S. latifolia*, for each sex-linked contig expression levels were estimated using read counts from both X/Y and X-hemizygous informative SNPs. SNPs were attributed to an X/Y or X hemizygous segregation type if the according posterior probability was higher than 0.5. SNPs are considered informative if the father is heterozygous and has a genotype that is different from the mother (otherwise it is not possible to tell apart the X from the Y allele and therefore it is not possible to compute X and Y expression separately). X/Y SNPs for which at least one female had over two percent of her reads belonging to the Y allele were removed as unlikely to be true X/Y SNPs. Informative autosomal SNPs from autosomal contigs were used in a similar way.

For contigs that only have X/X SNPs (SNPs for which the father's X is different to both Xs from the mother), Y expression level is only computed from the father as all males are homozygous in the progeny. Such contigs were therefore removed when having under 3 X/X SNPs to avoid approximations on the contig mean Y/X expression level (39 contigs removed in the flower buds dataset, 44 in the leaves dataset and 40 in the seedlings dataset).

In order to make *S. latifolia* expression levels comparable to *S. viscosa* and *S. vulgaris* for sex-linked contigs, *S. vicosa* and *S. vulgaris* expression levels were estimated using only the positions used in *S. latifolia* (informative X/Y or X-hemizygous SNPs). The read count of every position in every contig and for every *S. viscosa* and *S. vulgaris* individual was given by GATK DepthOfCoverage^15^. Only positions corresponding to informative autosomal, X/Y or X-hemizygous SNPs in *S. latifolia* were used to compute the expression level for each contig and each individual as explained in equation (1).

Contigwise *S. latifolia* autosomal, X, Y, X+X, X+Y allelic expression levels were then averaged among individuals. Autosomal normalised expression levels in the two outgroups (S. *vulgaris* and *S. viscosa)* were averaged together.

### 7) Identification of contigs with sex-biased expression

The analysis was done separately for the three tissues (flower buds, seedling and rosette leaves) as in Zemp et al.^8^ using the R package edgeR^27^. See Supplementary Table S2 for number of sex-biased contigs removed in order to study dosage compensation. Male-limited expressed contigs were identified by calculating the mean expression values (FPKM) in both sexes and selecting those which were exclusively expressed in males.

## Supplementary Figures

**Supplementary Figure S1:**
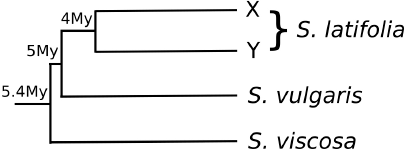
Relatedness among the three studied species, extracted from^31^ ages at the nodes are shown in million years (My). The exact relationship among species is poorly resolved^31–33^. In some phylogenies *S. viscosa* is closest to *S. latifolia*, whereas in others *S. vulgaris* is closest as shown here, and in others both species are equally diverged to *S. latifolia.*

**Supplementary Figure S2:**
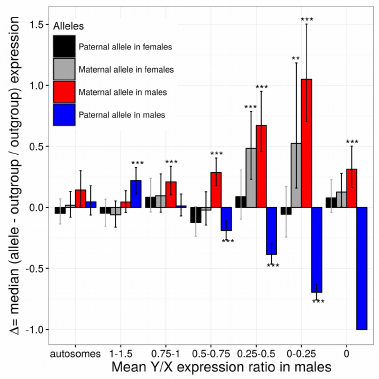
Normalised difference in allelic expression levels between *S. latifolia* and the two outgroups without sex chromosomes *S. vulgaris* and *S. viscosa* (hereafter Δ), in autosomal and sex-linked contigs for the **seedling** tissue. Maternal and paternal allelic read numbers were summed at SNP positions and normalised for each individual separately, then averaged among individuals for each contig. Δ was computed as follows: Δ=(allelic expression in *S. latifolia* – allelic expression in the outgroup) / allelic expression in the outgroup). If Δ is lower, higher or equal to zero, then expression in *S. latifolia* is respectively lower, higher or equal to the outgroup. For all contig categories, Δ was compared to zero using a Wilcoxon test. The median Δ, confidence intervals and p-values adjusted for multiple testing using a Benjamini and Hochberg correction are shown (***: p-value < 0.001; **: p-value < 0.01, *: p-value < 0.05). The Y/X ratio was computed in S. *latifolia* males and averaged among individuals to use as a proxy for Y degeneration. X-hemizygous contigs have a Y/X ratio equal to zero. Contigs with sex-biased expression were removed, as well as contigs with Y/X expression ratios above 1.5. Sample sizes for the different contig categories are: autosomal:200; 1-1.5:148; 0.75-1:139; 0.5-0.75:160; 0.25-0.5:114; 0-0.25:79; 0:205 (note that 200 autosomal contigs were randomly selected in order to have similar statistical power among gene categories). In the absence of dosage compensation, the single X in males should be expressed at levels similar to the outgroup that does not have sex chromosomes, in other words, without dosage compensation Δ should be close to zero for the maternal allele in males (red bars). Results show that the maternal allele is hyper-expressed in *S. latifolia* when the Y chromosome is degenerated, both in males and females.

**Supplementary Figure S3:**
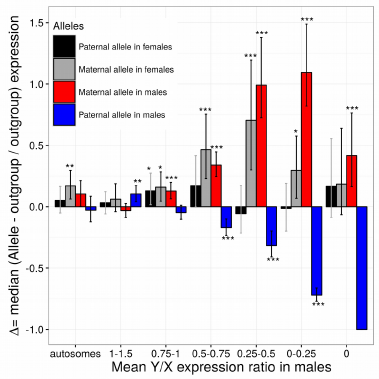
Normalised difference in allelic expression levels between *S. latifolia* and the two outgroups without sex chromosomes *S. vulgaris* and *S. viscosa* (Δ), in autosomal and sex-linked contigs for the **flower bud** tissue. Same legend as Supplementary Figure S2 except for sample sizes for the different contig categories: autosomal:200; 1-1.5:95; 0.75-1:195; 0.5-0.75:203; 0.25-0.5:176; 0-0.25:116; 0:103.

**Supplementary Figure S4:**
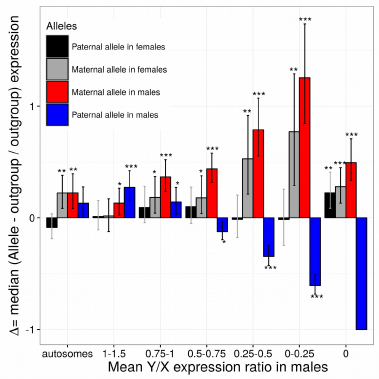
Normalised difference in allelic expression levels between *S. latifolia* and the two outgroups without sex chromosomes *S. vulgaris* and *S. viscosa* (Δ), in autosomal and sex-linked contigs for the **leaf** tissue. Same legend as Supplementary Figure S2 except for sample sizes for the different contig categories: autosomal:200; 1-1.5:159; 0.75-1:132; 0.5-0.75:147; 0.25-0.5:126; 0-0.25:71; 0:275.

**Supplementary Figure S5:**
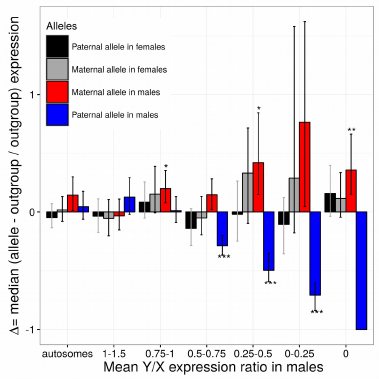
Normalised difference in allelic expression levels between *S. latifolia* and the two outgroups without sex chromosomes *S. vulgaris* and *S. viscosa* (Δ), in autosomal and sex-linked contigs that were **validated** (see Materials and Methods), for the **seedling** tissue. Same legend as Supplementary Figure S2 except for sample sizes for the different contig categories: autosomal:77; 1-1.5:71; 0.75-1:82; 0.5-0.75:91; 0.25-0.5:44; 0-0.25:29; 0:89.

**Supplementary Figure S6:**
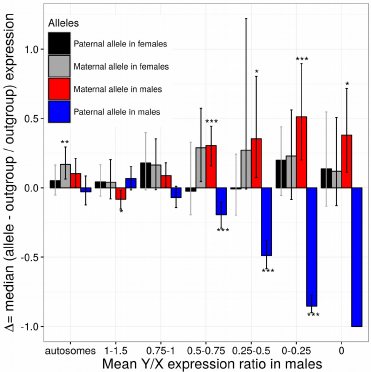
Normalised difference in allelic expression levels between *S. latifolia* and the two outgroups without sex chromosomes *S. vulgaris* and *S. viscosa* (Δ), in autosomal and sex-linked contigs that were **validated** (see Materials and Methods), for the **flower bud** tissue. Same legend as Supplementary Figure S2 except for sample sizes for the different contig categories: autosomal:74; 1-1.5:86; 0.75-1:91; 0.5-0.75:67; 0.25-0.5:45; 0-0.25:31; 0:55.

**Supplementary Figure S7:**
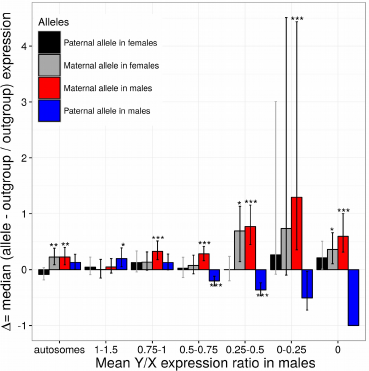
Normalised difference in allelic expression levels between *S. latifolia* and the two outgroups without sex chromosomes *S. vulgaris* and *S. viscosa* (Δ), in autosomal and sex-linked contigs that were **validated** (see Materials and Methods), for the **leaf** tissue. Same legend as Supplementary Figure S2 except for sample sizes for the different contig categories: autosomal:79; 1-1.5:84; 0.75-1:74; 0.5-0.75:77; 0.25-0.5:52; 0-0.25:19; 0:119.

**Supplementary Figure S8:**
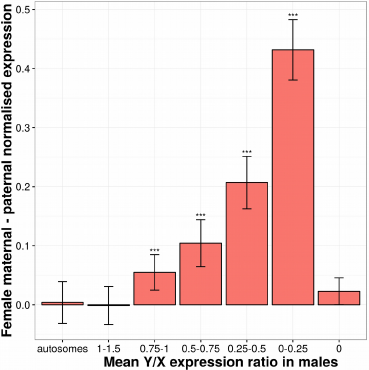
Normalised expression difference between the maternal and paternal allele in *S. latifolia* females in autosomal and sex-linked contigs for the **seedling** tissue. The Y axis unit is the normalised allelic read count difference in log scale. A linear regression model with mixed effects was used to study allelic expression in *S. latifolia* for every SNP position. In order to measure the changes in *S. latifolia* expression due to sex chromosomes evolution, the outgroup *S. vulgaris* that does not have sex chromosomes was used as a reference in the model (see Materials and Methods for details). The framework provided estimates for the normalised difference between the effect of paternal and maternal origin of alleles in interaction with the contig status (autosomal or sex-linked with various levels of Y degeneration), while accounting for inter-contig and inter-individual variability. See Supplementary Figure S2 legend for sample sizes for the different contig categories and statistical significance symbols. Results show that Y degeneration is linked to a significant expression difference between the paternal and maternal alleles in females, which is not observed in autosomal and non-degenerated sex-linked contigs.

**Supplementary Figure S9:**
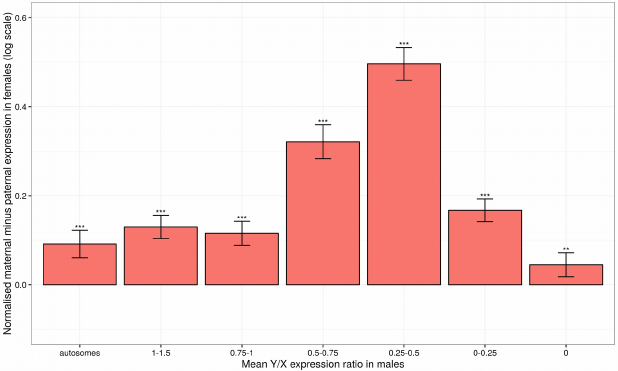
Normalised expression difference between the maternal and paternal allele in *S. latifolia* females in autosomal and sex-linked contigs for the **flower bud** tissue. See supplementary Figure S8 for legend and Supplementary Figure S3 for sample sizes for the different contig categories.

**Supplementary Figure S10:**
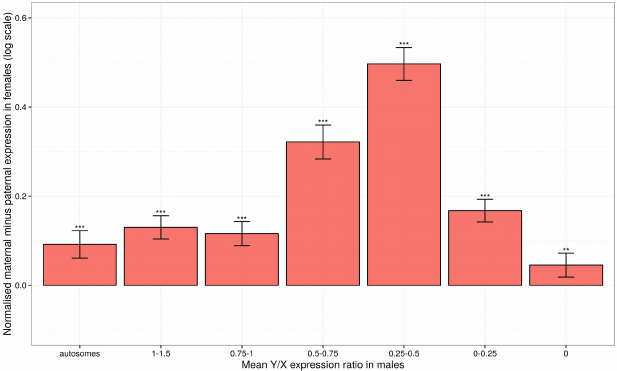
Normalised expression difference between the maternal and paternal allele in *S. latifolia* females in autosomal and sex-linked contigs for the **leaf** tissue. See supplementary Figure S8 for legend and Supplementary Figure S4 for sample sizes for the different contig categories.

**Supplementary Figure S11:**
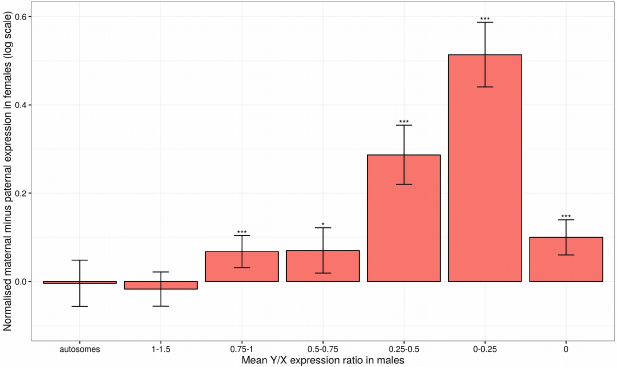
Normalised expression difference between the maternal and paternal allele in *S. latifolia* females in autosomal and sex-linked **validated** contigs for the **seedling** tissue. See supplementary Figure S8 for legend and Supplementary Figure S5 for sample sizes for the different contig categories.

**Supplementary Figure S12:**
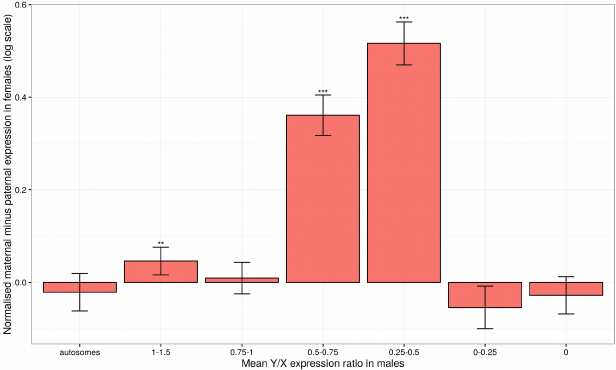
Normalised expression difference between the maternal and paternal allele in *S. latifolia* females in autosomal and sex-linked **validated** contigs for the **flower bud** tissue. See supplementary Figure S8 for legend and Supplementary Figure S6 for sample sizes for the different contig categories.

**Supplementary Figure S13:**
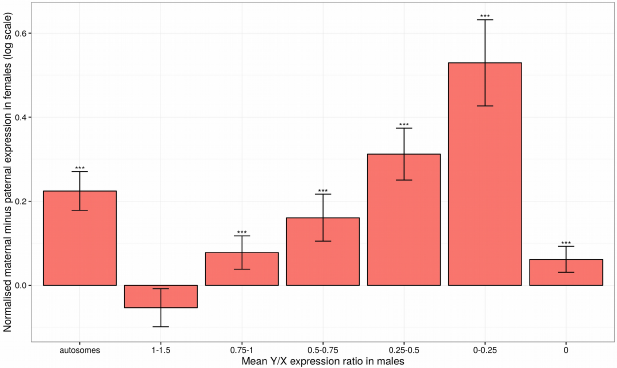
Normalised expression difference between the maternal and paternal allele in *S. latifolia* females in autosomal and sex-linked **validated** contigs for the **leaf** tissue. See supplementary Figure S8 for legend and Supplementary Figure S7 for sample sizes for the different contig categories.

**Supplementary Figure S14:**
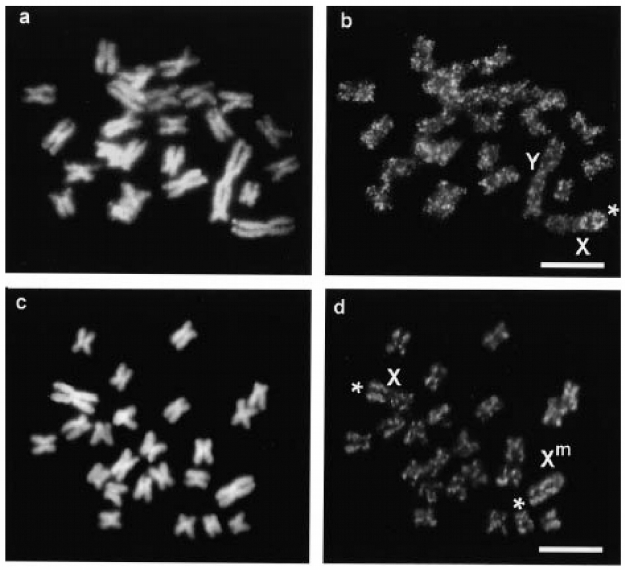
Original DNA methylation staining results from Siroky et al 1998^34^. **(a)** Male metaphase chromosomes stained with PI. **(b)** FITC-anti-5-mC signals on the same chromosomes. The hypomethylated shorter X arm is marked by an asterisk; The X and Y chromosomes are indicated. **(c)** Female metaphase chromosomes stained with PI. **(d)** FITC-anti-5-mC signals on the same chromosomes. Shorter arms of the Xs are indicated by asterisks. The hypermethylated X chromosome is marked as X^m^. Bars = 5μm.

**Supplementary Table S1:** library sizes (number of reads) of each individual and mapping statistics.

**Supplementary Table S2:** Number of contigs after SEX-DETector inferences, removal of sex-bias and selection of validated contigs in the three tissues.

|  | Tissue type |  |  |
| --- | --- | --- | --- |
|  | flower buds | leaves | seedlings |
| number of ORFs | 46178 |  |  |
| Unassigned | 33172 | 33564 | 33781 |
| Autosomal | 11662 | 11558 | 11292 |
| X/Y | 1140 | 772 | 844 |
| X-hemizygous | 204 | 284 | 261 |
| X/Y non sex-biased | 901 | 733 | 732 |
| X-hemizygous non sex-biased | 103 | 275 | 205 |
| X/Y non sex-biased validated | 339 | 345 | 365 |
| X-hemizygous non sex-biased validated | 55 | 119 | 89 |
| Autosomal validated | 74 | 79 | 77 |

Supplementary Table S3: list of known sex-linked genes in *S. latifolia* and associated literature references.

